# Full-length LINE-1s with functional ORFs are favoured by host-level selection in human genomes

**DOI:** 10.1101/2020.01.09.900191

**Authors:** Heinrich zu Dohna, Bo Zhou, Xiaowei Zhu, Alexander E. Urban

**Affiliations:** Department of Biology, American University of Beirut, 11–0236, Beirut, Lebanon; Departments of Psychiatry and Genetics, Stanford University School of Medicine, Stanford, CA 94305, USA

**Author notes:** email: Heinrich Dohna; Bo Zhou; Xiaowei Zhu; Alexander E Urban.

## Abstract

The effects of transposable elements, such as LINE-1, on host fitness are still poorly understood. Our analysis of the site frequency spectrum of LINE-1s and SNPs within LINE1s in human genomes shows that selection coefficients are higher for full-length than for fragment LINE-1s and on full-length LINE-1s, SNPs are significantly depleted on ORFs and non-synonymous sites within ORFs. These results suggest that host-level selection maintains transposition competent LINE-1s.

## Main

LINE-1 (long interspersed element 1) retrotransposons are mobile genetic elements that play an important role in modifying the human genome ^1^. LINE-1 insertions are generally considered disruptive since they are depleted in regulatory regions ^2,3^ and are associated with genetic disorders and diseases ^4^. The destabilizing effects of LINE-1 insertions led to the evolution of LINE-1 silencing mechanisms, such as LINE-1 methylation ^5^ and piRNAs ^6^.

However, there is also evidence that retrotransposition has become an integral part of the host developmental program ^7,8^. Yet, so far, there is no direct evidence that LINE-1 retrotransposition is beneficial to the human host. In this study, we used two methods to analyze signals of host-level selection on retrotransposition activity. We estimated the selection coefficients of polymorphic LINE-1s from the site frequency spectrum (SFS) of LINE-1 insertions in the human population and we analyzed patterns of single-nucleotide polymorphism (SNP) depletion on LINE-1 insertions within the human reference genome hg19.

We obtained the LINE-1 SFS from Gardner et al. ^9^ and fitted models to this SFS in which the selection coefficient of a LINE-1 insertion could vary by LINE-1 insertion length and between full-length and fragment LINE-1s, following the approach by Boissinot et al. ^10^. According to the best-fitting model (i.e., the model with the lowest AIC), LINE-1 selection coefficients decreased with LINE-1 length but increased for full-length LINE-1 (Fig. 1a; Supplementary Table 1). LINE-1s of all lengths had a negative selection coefficient, but short fragments and full-length LINE-1s had the highest estimated selection coefficients (Fig. 1a).

**Figure 1:**
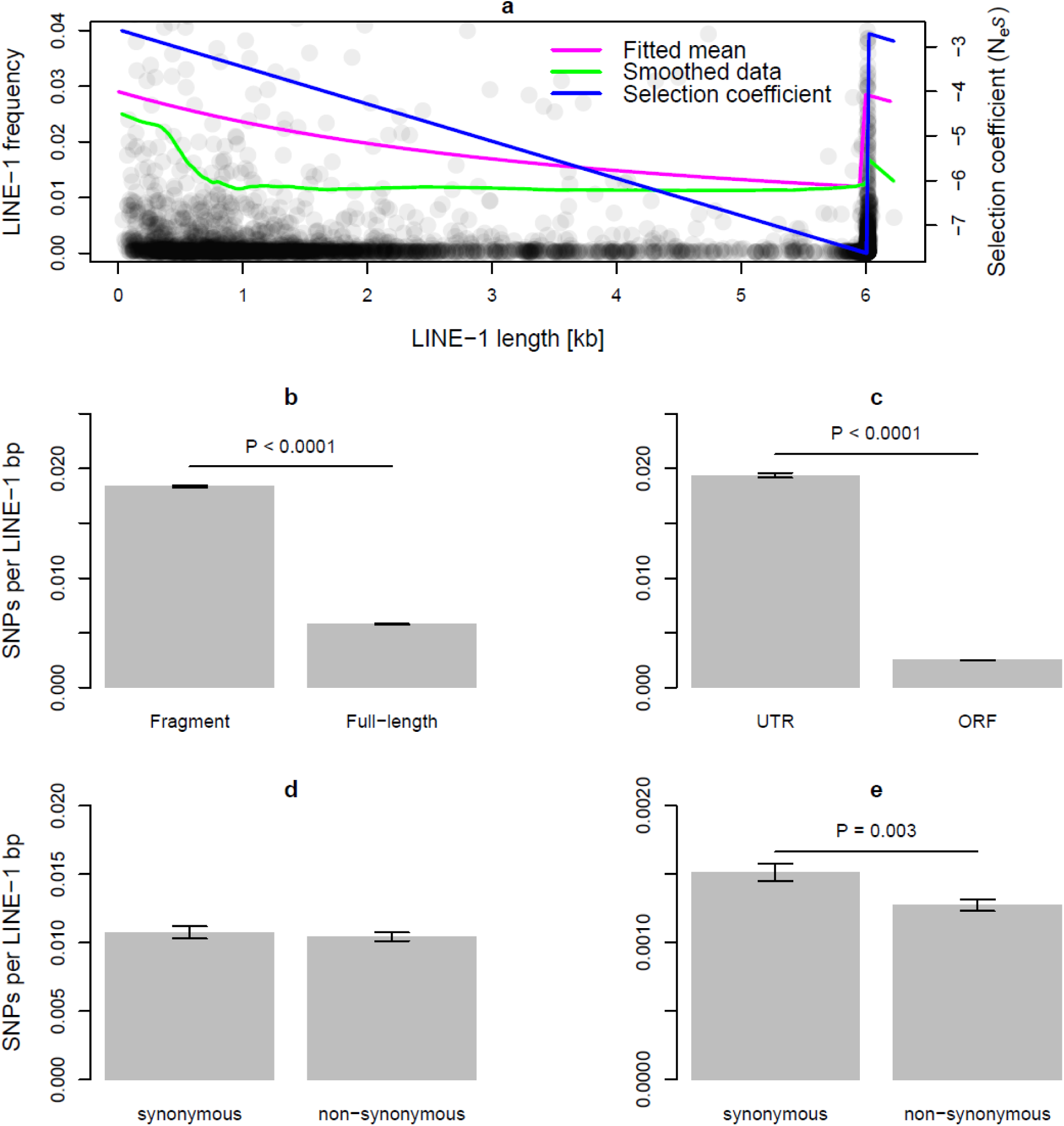
(a) LINE-1 frequency vs. LINE-1 length. Points show frequency and lengths of individual LINE-1. (The frequency range from zero to 0.04 was chosen to illustrate the patterns best; 7% of the frequency values were above 0.04 and are therefore not shown on the plot.) The green curve shows a smoothing spline fitted through the points, magenta curve shows the predicted average frequency per length based on best-fitting model, and blue line shows selection coefficient as function of LINE-1 length for best-fitting model. (b-e) Number of polymorphic positions per LINE-1 bp. (b) Comparison between fragment and full-length LINE-1s, (c) between ORFs and UTRs on full-length LINE-1s, (d) between synonymous and non-synonymous positions on full ORFs on fragment LINE-1s, and (e) between synonymous and non-synonymous positions on full ORFs on full-length LINE-1s. *P*-values were determined by logistic regression.

One potential limitation of our SFS analysis is the possibility of spurious correlations between LINE-1 insertion length and frequency due to frequency-dependent length estimates. We applied the mobile element detection tool MELT ^9^ to simulated reads and found that the accuracy of the estimated LINE-1 insertion length does indeed depend on LINE-1 frequency. However, when we averaged over uncertainties in length estimation (see Material and Method section *Uncertainties in LINE-1 detection and length estimation*) the results were qualitatively the same as above (Supplementary Table 1).

Another limitation of our SFS analysis was that it assumed a constant population size, a mutation-selection-drift equilibrium, and ignored background selection and population structure. However, any estimation bias due to these assumptions is expected to affect LINE-1 insertions of different lengths in a similar way and should therefore not affect inferences about the relationship between selection coefficient and LINE-1 length. In line with this assertion, we observed an excess of observed low-frequency LINE-1s compared to model predictions that was consistent across different LINE-1 length classes (Supplementary Fig. 1). While the excess of low frequency LINE-1 indicates that the model we fitted to the data did not capture some aspects of the mutation-selection process, the consistency across LINE-1 length classes showed that our model simplifications were unlikely to affect the inferred relationship between LINE-1 length and selection coefficient.

The decrease of LINE-1 selection coefficients with insertion length that our analysis uncovered is most likely due to increased ectopic recombination between longer LINE-1 insertions^11,12^. The increased selection coefficient for full-length LINE-1s suggests a beneficial effect of retrotransposition for the host since only full-length LINE-1s are retrotransposition competent. It is important to note that detecting the higher selection coefficient for full-length LINE-1s required accounting for the decrease of selection coefficient with LINE-1 length by fitting an appropriate model to a dataset large enough to include the relatively rare long LINE-1 fragments. Previous analyses of LINE-1 selection coefficients failed to account for the decrease of selection coefficient with LINE-1 length, either because of a small sample size ^10^ or by fitting a selection model without explicit length-dependency ^13^, and therefore missed that full-length LINE-1s have a significantly higher selection coefficient than long fragments.

We also analyzed how SNPs from the 1000 Genome dataset are distributed within LINE-1 insertions. The probability of a base pair within a LINE-1 containing a SNP depended significantly on the immediate tri-nucleotide neighborhood, increased with the number of SNPs in regions flanking the LINE-1, decreased with read coverage, and increased with the proportion of nucleotides that differed from the human LINE-1 (L1HS) consensus (Supplementary Table 2). More importantly, the probability for a SNP was significantly lower for full-length than fragment LINE-1s, for coding regions, and for non-synonymous sites within coding regions (odds ratios = 0.78, 0.63, and 0.76, respectively, all P < 0.0001, Supplementary Table 2, Fig. 1b,c,e). The reduction in SNP probability on coding regions and on non-synonymous positions within coding regions was significantly more pronounced on full-length than fragment LINE-1s (odds ratio for SNPs on coding vs. non-coding region was 0.63 on fragment LINE-1s and 0.26 on full-length LINE-1, Supplementary Table 2, Fig. 1 c). When the analysis was restricted to LINE-1 ORFs without frameshifts, fragment LINE-1s showed no SNP reduction on non-synonymous sites (Fig. 1d), whereas on full-length LINE-1s the SNP density was significantly reduced on non-synonymous sites (odds ratio = 0.84, P = 0.003, Fig. 1e). All results were qualitatively the same, regardless of whether or not SNPs with a significant excess of heterozygotes and a quality score below 100 were excluded from the analysis.

The reduction of SNPs on full-length LINE-1s compared to fragments could be explained by transcription-coupled repair ^14^. However, the reduction of SNPs on ORF1 and ORF2 and in particular the additional reduction of SNPs on non-synonymous sites cannot be explained easily by differential mutation, transcription-coupled DNA repair, or SNP detection but instead suggests that purifying selection removes single-nucleotide mutations that alter the translated amino acid sequence. It is important to note that our analysis concerned SNPs that accumulate post-LINE-1 insertions. These SNPs are influenced by selection on the host level rather than transposon level. Our results are therefore a strong indication that retrotransposition fulfills a function for the host that is maintained by selection on the host level.

It is unclear why functional full-length LINE-1s would be beneficial for the host. There are several lines of evidence that retrotransposition is damaging for the host, such as the co-evolutionary arms race between LINE-1s and LINE-1-silencing KRAB zinc fingers ^15^, the negative correlation between LINE-1 retrotransposition activity and population frequency ^16^, and the negative selection coefficients of LINE-1s as shown by our and previous results ^10^. The evidence for detrimental effects of LINE-1 insertions appears in conflict with our results. However, all evolutionary evidence for detrimental effects of LINE-1 insertions concerns germline insertions. Hence, we hypothesize that germline LINE-1 insertions are generally detrimental but host-level selection maintains the retrotransposition machinery due to beneficial effects of somatic LINE-1s insertions. The most likely candidates for such beneficial effects are somatic LINE-1 insertions in neurons^8^. Identifying what aspect of the LINE-1 retrotransposition is beneficial for the host will be a major step in our understanding of LINE-1 host co-evolution in humans and other mammals.

## Methods

### Frequencies of LINE-1 insertions and deletions in 1000 Genome data

We downloaded from dbVar vcf files with genomic locations and population-level frequencies of non-reference LINE-1 insertions in genomes of the 1000 Genomes Project Phase III (ftp://ftp.ncbi.nlm.nih.gov/pub/dbVar/data/Homo_sapiens/by_study/vcf/nstd144.GRCh37.variant_region.vcf). In addition, we used MELT ver. 2.0 (http://melt.igs.umaryland.edu) to identify deletions of reference LINE-1s on the human genome in the high coverage genomes (30X) of the 1000 Genomes Project.

### Uncertainties in LINE-1 detection and length estimation

We simulated 50 genomes with known LINE-1 insertions using the read simulator wgsim ^17^. We selected 50 samples from the 1000 Genomes data and collected for each sample the information about genotype, genomic location, start, and endpoint of all non-reference LINE-1 insertions that were called in the 1000 Genomes data. Next, we generated two whole-genome sequences that included the called insertions and generated haploid reads for each sequence using wgsim, and combined these reads to simulate a diploid genome with the same coverage as the 1000 Genome samples. We then ran MELT ver. 2.0 with the “Group” option on these simulated genomes. Eleven of the 50 simulated genomes produced an error message when analyzed by MELT. We estimated from the remaining 39 genomes via logistic regression the detection probability of LINE-1s as a function of LINE-1 length, LINE-1 frequency, and an indicator for full-length LINE-1. We furthermore grouped the true and estimated LINE-1 lengths into bins of 500 bp width and estimated a discretized joint probability of true and estimated LINE-1 length. We fitted a logistic regression to determine how the probability of the true and estimated LINE-1 length being in the same bin changes as a function of LINE-1 frequency.

### Signals of LINE-1 selection in 1000 Genome data

We analyzed whether selection coefficients change with LINE-1 insertion length and differ between full-length and fragment LINE-1s. Specifically, we tested the influence of two predictor variables, namely LINE-1 length and an indicator for LINE-1 length being above 6000 bp (i.e. indicator for full-length), on the selection coefficient *s*, following the approach outlined by Boissinot et al. ^10^. This approach assumes that a LINE-1 with selection coefficient *s* confers, respectively, fitness values 1+*s*, 1+*s*/2 and 1 to individuals who are homozygous carriers of the LINE-1, heterozygous, or without the LINE-1 and derives an SFS, assuming a mutation-selection-drift balance. We slightly modified the approach outlined in ref. ^10^ by including a probability that a LINE-1 is present on the reference genome as function of LINE-1 frequency, estimating this probability from 1000 Genome data via logistic regression, introducing detection sensitivity *p* (obtained from ref. ^9^) and calculating the probability of an inclusion of a non-reference LINE-1 in the study as 1 – (1 – *px*)^*n*^, where *p* stands for sensitivity, *x* for the allele frequency and *n* for the number of samples. We estimated via maximum likelihood the selection coefficients of LINE-1s from the LINE-1 SFS, assuming a population size of 10^5^. We fitted four different models for the selection coefficient *s* of LINE-1s. In the simplest model, the coefficient *s* was the same for all LINE-1s. In the other three models, *s* was a linear function of either one or both of the two predictor variables, LINE-1 length and the indicator for full-length LINE-1s. We compared the Akaike Information Criterion (AIC) between the four models and determined the best-fitting model as the one with the lowest AIC. We ran this analysis, either treating estimated LINE-1 length values as observations, or incorporating the frequency-dependent uncertainty in LINE-1 length estimation. To account for uncertainty in LINE-1 length estimation we used the discretized joint probability of true and estimated LINE-1 insertion length as estimated from the simulation study and calculated a likelihood of each LINE-1 insertion as

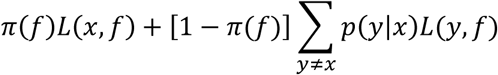

Where *x* and *y* denote the estimated and true length of a LINE-1 insertion, respectively, *π*(*f*) denotes the probability that the estimated and true length of a LINE-1 insertion are in the same length bin, *f* denotes the population frequency of a LINE-1, *L*(*x*,*f*) the likelihood of a LINE-1 insertion of length *x* at frequency *f*, and *p*(*y*|*x*) the probability of true length *y* given the estimated length *x*, conditional on *x* ≠ *y*.

### SNP density in full-length and fragment LINE-1

We estimated via logistic regression the effect of various predictors on SNP density in reference LINE-1s. Each base pair within L1HS on hg19 was counted as an observation, and the response variable was whether or not a base pair contained a SNP within the 1000 Genome data. We included for each base pair the following covariates: (i) the trinucleotide surrounding a base pair and (ii) the number of SNPs within 1000 bp flanking the LINE-1 on either side to control for variation in the mutation rate due to immediate and wider sequence context; (iii) the proportion of nucleotides within the same LINE-1 that differed from the L1HS consensus (obtained from repbase https://www.girinst.org/repbase/) to control for an accumulation of SNPs due to age of a LINE-1 insertion; (iv) the average coverage on a base pair among all 1000 Genomes samples to control for variation in SNP detectability; (v) whether a LINE-1 insertion was inside a promoter region, exon, or intron. Our regression model also included the following three predictor variables to capture effects of selection on LINE-1 retrotransposition capacity: (i) whether or not the LINE-1 was full-length (above 6000 bp length), (ii) whether or not the base-pair was on a coding sequence within a LINE-1 (ORF1 or ORF2) and (iii) whether or not a base-pair was on a non-synonymous position within a LINE-1 ORF. We treated the first two base-pairs of each codon within a coding sequence as non-synonymous positions. We used the first and last nine nucleotides of each ORF on the L1HS consensus sequence as motifs to identify beginning and end of ORFs within LINE-1s. We first searched for these, allowing for one mismatch for the start motifs and two mismatches for the end motifs. After identifying the start and end motifs of each ORF we counted forward from the start codon and backward from the stop codon to identify codon start positions and formed for each ORF an intersection of these positions to exclude frameshifts. We ran an additional analysis with the same set of predictor variables but using only bps with full-length ORFs without frameshifts.

### SNPs due to mismapped LINE-1 reads

Because LINE-1 sequences are highly repetitive in the genome there is a possibility that some SNPs on reference LINE-1s in the 1000 Genome data could have been due to mismapping reads. Since a locus with mismapped reads is expected to also contain reads from that location, mismapping should produce the signal of a heterozygous genotype. We therefore repeated the analysis above, excluding any SNP that showed a significant deviation from Hardy-Weinberg genotype frequencies in any of the five populations and an excess of heterozygote genotypes.

## Author contributions

H.D. and A.U. conceived the study. H.D. wrote the manuscript. All authors read and commented on the manuscript. B.Z. and X.Z. provided expertise and crucial input for study design and data analysis. H.D. performed the data analysis.

