## Supplementary Information for "Full-length LINE-1s with functional ORFs are favoured by host-level selection in human genomes"

*Corresponding author

### Supplementary Information

Table 1: Models for selection coefficient fitted to LINE-1 SFS and length estimates in 1000 Genome data

| Predictors of the selection coefficient | Parameter values | Parameter values* | AIC | ΔAIC | AIC* | ΔAIC* |
|---|---|---|---|---|---|---|
| none | $\beta_0$ = -3.4*$10^{-5}$ | $\beta_0$ = -5.2*$10^{-5}$ | 30607 | 54 | 29220 | 83 |
| L1 length ($\beta_1$) | $\beta_0$ = -3.4*$10^{-5}$<br>$\beta_1$ = -2.1*$10^{-10}$ | $\beta_0$ = -5.4*$10^{-5}$<br>$\beta_1$ = 9.5*$10^{-10}$ | 30609 | 56 | 29221 | 84 |
| L1 full-length indicator ($\beta_1$) | $\beta_0$ = -3.7*$10^{-5}$<br>$\beta_1$ = 1.0*$10^{-5}$ | $\beta_0$ = -6.0*$10^{-5}$<br>$\beta_1$ = 2.5*$10^{-5}$ | 30597 | 44 | 29191 | 53 |
| L1 length ($\beta_1$) and full-length indicator ($\beta_2$) | $\beta_0$ = -2.6*$10^{-5}$<br>$\beta_1$ = -8.3*$10^{-9}$<br>$\beta_2$ = 4.9*$10^{-5}$ | $\beta_0$ = -3.8*$10^{-5}$<br>$\beta_1$ = -1.5*$10^{-8}$<br>$\beta_2$ = 9.8*$10^{-5}$ | 30553 | 0 | 29138 | 0 |

*estimates obtained by incorporating uncertainties in LINE-1 length estimation

Table 2: Full set of predictors of SNP probability

| Predictors of SNP probability | Coefficient | *Exp*(Coefficient) | P-value |
|---|---|---|---|
| (Intercept) | -5.73 | 0.003 | <0.0001 |
| Trinucleotide AAC | 0.33 | 1.39 | <0.0001 |
| Trinucleotide AAG | 0.13 | 1.14 | 0.007 |
| Trinucleotide AAT | 0.6 | 1.83 | <0.0001 |
| Trinucleotide ACA | 0.79 | 2.2 | <0.0001 |
| Trinucleotide ACC | 1.07 | 2.91 | <0.0001 |
| Trinucleotide ACG | 3.22 | 25.09 | <0.0001 |
| Trinucleotide ACT | 0.84 | 2.32 | <0.0001 |
| Trinucleotide AGA | 0.64 | 1.89 | <0.0001 |
| Trinucleotide AGC | 0.84 | 2.31 | <0.0001 |
| Trinucleotide AGG | 0.84 | 2.31 | <0.0001 |
| Trinucleotide ATA | 0.68 | 1.96 | <0.0001 |
| Trinucleotide ATC | 0.28 | 1.32 | <0.0001 |
| Trinucleotide ATG | 0.87 | 2.39 | <0.0001 |
| Trinucleotide CAA | 0.27 | 1.31 | <0.0001 |
| Trinucleotide CAC | 0.48 | 1.61 | <0.0001 |
| Trinucleotide CAG | 0.39 | 1.48 | <0.0001 |
| Trinucleotide CCA | 0.6 | 1.82 | <0.0001 |
| Trinucleotide CCC | 0.85 | 2.33 | <0.0001 |
| Trinucleotide CCG | 3.27 | 26.32 | <0.0001 |
| Trinucleotide CGA | 3.01 | 20.33 | <0.0001 |
| Trinucleotide CGC | 3.16 | 23.51 | <0.0001 |
| Trinucleotide CTA | 0.21 | 1.24 | <0.0001 |

| | | | |
|---|---|---|---|
| Trinucleotide CTC | 0.05 | 1.05 | 0.34 |
| Trinucleotide GAA | -0.12 | 0.89 | 0.015 |
| Trinucleotide GAC | 0.15 | 1.16 | 0.008 |
| Trinucleotide GCA | 0.62 | 1.85 | <0.0001 |
| Trinucleotide GCC | 0.99 | 2.7 | <0.0001 |
| Trinucleotide GGA | 0.71 | 2.03 | <0.0001 |
| Trinucleotide GTA | 0.2 | 1.22 | 0.0003 |
| Trinucleotide TAA | 0.27 | 1.32 | <0.0001 |
| Trinucleotide TCA | 0.48 | 1.61 | <0.0001 |
| SNP count on LINE-1 flanks | 0.02 | 1.02 | <0.0001 |
| Mean coverage | 0.01 | 1.01 | <0.0001 |
| LINE-1 length | 5.95*10-5 | 1 | <0.0001 |
| Proportion mismatch from consensus | 10.75 | 46700 | <0.0001 |
| Position in gene body | -0.07 | 0.94 | <0.0001 |
| Position on exon | 0.5 | 1.65 | <0.0001 |
| Position in promoter region | -0.11 | 0.9 | 0.02 |
| Full-length LINE-1 | -0.25 | 0.78 | <0.0001 |
| Non-synonymous position on LINE-1 | -0.28 | 0.76 | <0.0001 |
| Coding sequence on LINE-1 | -0.46 | 0.63 | <0.0001 |
| Interaction coding sequence on LINE-1 x full-length LINE-1 | -0.87 | 0.42 | <0.0001 |
| Interaction non-synonymous on LINE-1 x full-length LINE-1 | -0.74 | 0.48 | <0.0001 |

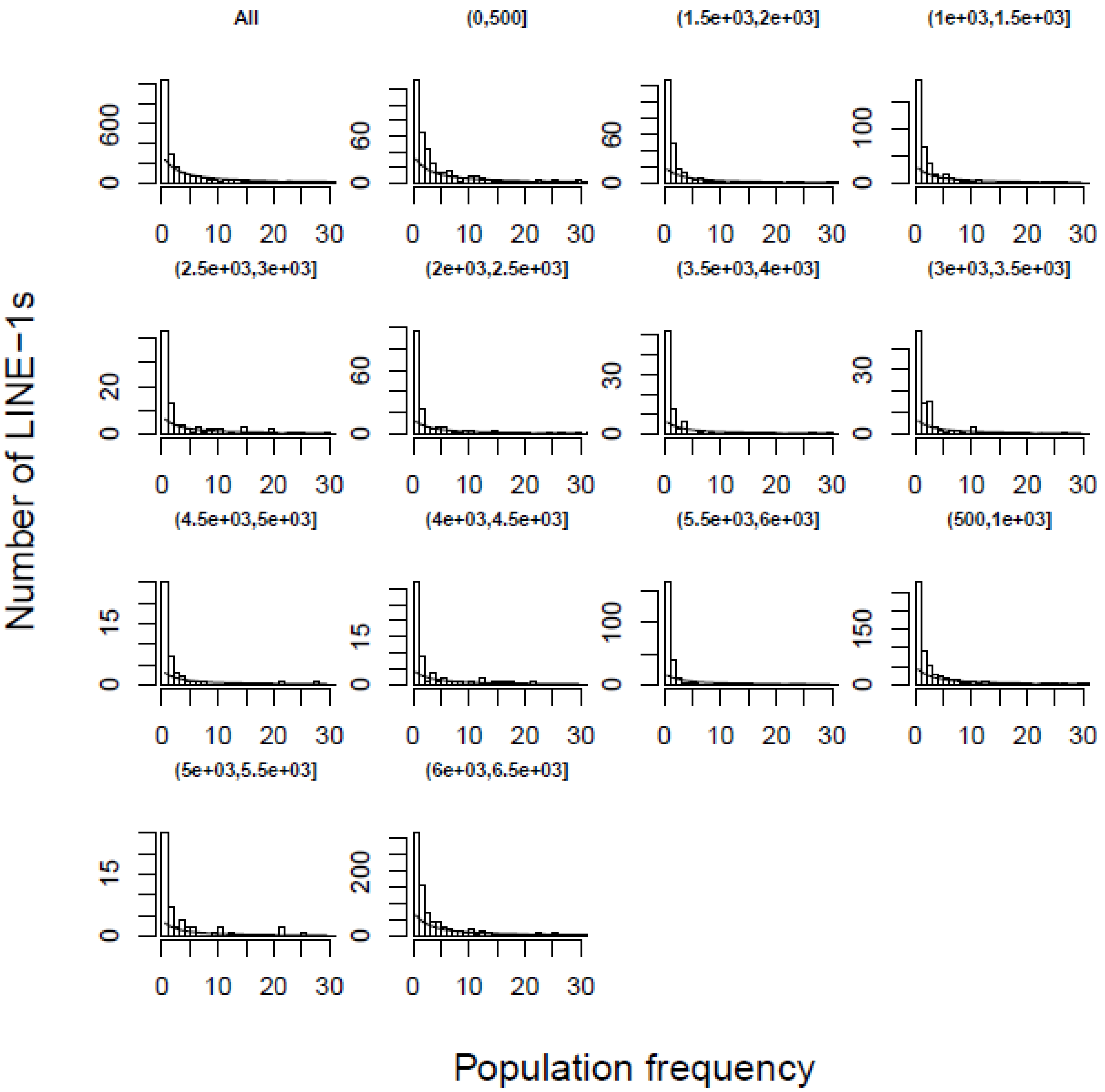


Figure 1: Observed and expected SFS. Bars indicate the observed number of LINE-1s that occur at a particular population frequency. Lines show the expected number according to the best-fitting model. The first panel shows the summary across all LINE-1 length classes. All other panels show the observed and expected numbers for the LINE-1 length class indicated by the interval in the panel title.
